# The effect of mating complexity on gene drive dynamics

**DOI:** 10.1101/2021.09.16.460618

**Authors:** Prateek Verma, R. Guy Reeves, Samson Simon, Mathias Otto, Chaitanya S. Gokhale

## Abstract

Gene drive technology promises to deliver on some of the global challenges humanity faces to-day in healthcare, agriculture and conservation. However, there is a limited understanding of the consequences of releasing self-perpetuating transgenic organisms into the wild populations under complex ecological conditions. In this study, we analyze the impact of three such complexities, mate-choice, mating systems and spatial mating network, on the population dynamics for two distinct classes of modification gene drive systems. All three factors had a high impact on the modelling outcome. First, we demonstrate that distortion based gene drives appear to be more robust against the mate-choice than viability-based gene drives. Second, we find that gene drive spread is much faster for higher degrees of polygamy. Including a fitness cost, the drive is fastest for intermediate levels of polygamy. Finally, the spread of gene drive is faster and more effective when the individuals have fewer connections in a spatial mating network. Our results highlight the need to include mating complexities while modelling the properties of gene drives such as release thresholds, timescales and population-level consequences. This inclusion will enable a more confident prediction of the dynamics of engineered gene drives and possibly even inform on the origin and evolution of natural gene drives.

## Introduction

Gene drive technology is being developed to potentially deliver on some of the critical challenges in human health, agriculture or biodiversity conservation (Brossard et al., 2019; Buchman et al., 2018; Johnson et al., 2016; Prowse et al., 2017; Windbichler et al., 2011). A prominent example of gene drive is to push transgenes into wild mosquito populations that make them resistant to malaria parasites (Carballar-Lejarazú et al., 2020; Gantz et al., 2015). In conservation studies, the potential of gene drives to control the spread of invasive species or implementing disease resistance in endangered species is being discussed (Godwin et al., 2019; Johnson et al., 2016; Prowse et al., 2017). In agriculture, gene drive could control pest populations like fruit flies in cherry plantations or transform the pest population to make them more susceptible to pesticides (Barrett et al., 2019; Buchman et al., 2018). To date, no gene drive organisms have been released into the wild populations. All gene drive constructs are necessarily transgenic and require the release of genetically modified organisms into wild populations. The possibility exists that not all the unintended consequences of gene drive releases are reversible. Consequently, modelling is key to evaluating this technology.

Theoretical and laboratory studies indicate that some transgenic driving constructs could spread through wild populations in a relatively small number of generations (Burt, 2003; Deredec et al., 2008; Simoni et al., 2020; Windbichler et al., 2011). However, such results may only be valid under ideal conditions, such as random mating and other simplified ecological interaction. Therefore, such estimates may not provide robust predictions of the drive’s behaviour under field conditions in some circumstances. Several studies related to the risk assessment of gene drives have highlighted the relevance of ecological and technological bottlenecks like resistance evolution, mate-choice, mating system, and spatial interaction in successfully deploying gene drive organisms (Collins, 2018; Giese et al., 2019; Moro et al., 2018; of Sciences Engineering and Medicine, 2016; Oye et al., 2014). Thus, assessing the validity of the model’s assumptions is an essential task that any gene drive technology needs to overcome to become an option for a field release. While numerous laboratory assumptions may be violated in the wild, we choose to focus on aspects relating to the ecological complexity of mating. We demonstrate that mate-choice, aspects of the mating system, and spatial aspects of finding mating partners can change the course of eco-evolutionary trajectories of gene drive systems.

Gene drive leverages sexual reproduction by biasing the inheritance of a specific gene from one generation to the next. Hence, it becomes imperative to account for the target species’ reproductive biology and mating pattern to predict critical parameters for a release, such as the threshold of gene-drive organisms (GDOs) needed in order to be successful (Moro et al., 2018; of Sciences Engineering and Medicine, 2016). While theoretical explorations and laboratory experiments often assume simplified mating conditions based on random mating, other factors also influence mating success in the wild. Non-random mating may result from various factors and processes, such as inbreeding, mate-choice, and multiple matings, often part of complex mating systems. These aspects have already been recognized in gene drive research (Deredec et al., 2008; Noble et al., 2017; Qureshi et al., 2019; Unckless et al., 2015). Inbreeding diminishing the frequency of heterozygotes in the population, slowing the spread of gene drive (Bull et al., 2019*b*; Champer et al., 2021). In natural meiotic drive, females of some species can discriminate against males carrying drive when the region containing the drive gene is linked to mate-choice signals (Price and Wedell, 2008; Wedell and Price, 2015). For example, the naturally occurring selfish genetic element (*t*-complex) in *Mus domesticus* exhibits mate preference whereby both sexes appear to avoid heterozygous mate using olfactory cues (Lenington, 1983, 1991; Lindholm et al., 2013).

A newly evolved natural distorter system may remain at low frequency due to reduced fertility of drive carrying individuals, with the resulting potential to selection for mating bias (Charlesworth and Charlesworth, 2010; Wedell and Price, 2015). However, it remains unclear if bias in mate preference can quickly evolve for laboratory-engineered gene drives. A study by Drury et al. (2017) showed that non-random mating caused by inbreeding could render the CRISPR based gene drive inefficient against standing genetic variation resulting in cleavage re-sistance for Cas9 target sites in the flour beetle *Tribolium castaneum*. Bull (2017) suggested that mild levels of initial inbreeding can lead to the evolution of selfing in hermaphrodites (plants) in response to a homing endonuclease gene drive. Suppression gene drives, aimed at the local eradication of target species, can lead to the evolution of sib-mating, significantly hampering the spread of the driven gene (Bull et al., 2019*b*).

The mating system of target species will also play an essential role in determining the population dynamics of the spread of gene drives. For example, even in the absence of pre-copulatory mate-choice, the *t*-haplotype meiotic drive in mice can be limited by their polyandrous mating system where females mate with multiple males in a breeding cycle (Lindholm et al., 2016; Manser et al., 2017). The *t*-haplotype carrying males have reduced fertility, so when a female mates with multiple males, the fertilization of non-drive carrying male due to sperm competition is more likely (Manser et al., 2020, 2017). A sex-linked gene drive based on utilizing *t*-haplotypes has been proposed to suppress the rodent populations (Godwin et al., 2019; Leitschuh et al., 2018). The impact of polyandry on the population-level dynamics of one such proposed gene drive construct (*t-Sry*) has been studied by Manser et al. (2019). *t-Sry* has two components: *t*-haplotypes and sex-determining *Sry* gene and polyandry negatively effect its spread Manser et al. (2019). Focusing on an age-structured population, Huang et al. (2009) showed that the mating system for Medea and engineered under-dominance gene drives can significantly change the predicted threshold number of released transgenic individuals for successful population transformation. They also found that low polyandry levels can hamper gene drive spread if only males are released. When the gene drive causes male scarcity (Y-shredder), in polygamous systems where males mate with multiple females hampers, the gene spread efficacy (Prowse et al., 2019).

Most wild populations do not exist in a single panmictic population but often as multiple heterogeneous communities. In a spatially segregated population, individuals are more likely to interact with others in their vicinity than randomly with everyone in the population. Some mathematical models of gene drive use numerical evaluation in continuous space to account for such spatial interaction (Beaghton et al., 2017; Girardin et al., 2019; Tanaka et al., 2017). In these systems, the time required for a gene to spread depends on the interaction zone where the wildtype meets the transgenics. This zone is the wave’s leading edge in the reaction-diffusion models (Beaghton et al., 2017; Girardin et al., 2019; Tanaka et al., 2017). In the case of suppression drives, the wave sweeps through the wild population, leaving empty space (Barton and Turelli, 2011; Bull et al., 2019*a*; North et al., 2013). Compared to the panmictic models, the suppression drive can be less effective and slow in spatial models (Champer et al., 2021, 2020; North et al., 2013). When considering long-range dispersal, the wild-types could occupy the empty space created by the suppression drive resulting in local cycles of drive eradication and reoccupation by the wild-type (Champer et al., 2021). Similar cyclical dynamics is possible for reversal drives released to convert the previously established homing drives (Girardin et al., 2019). A question primarily ignored in some of these spatial models concerns the effect of heterogeneous interaction among individuals during mating. For example, the interactions in mathematical models using reaction-diffusion equations are assumed to be homogeneous. The spread of the gene drives relies on sexual reproduction, which is most likely not spatially or temporally uniform for all individuals in a population. A population structured on a network can help account for the natural heterogeneity in mating success. We use concepts from network theory and build a model to investigate how spatial mating networks could affect the gene drive’s spread.

Risk assessors face fundamental challenges when using models in their assessments. First, understanding modelling approaches and the underlying assumptions for complex applications like synthetic gene drives is far from trivial. Second, evaluating the effects of ecological factors on gene drive efficacy is not intuitive. Hence, in general, risk assessment of GDOs will be complex and include more uncertainties than current GM crops designed for release into the environment (Simon et al., 2018). Analogous to other risk assessments (EFSA document on good modelling practice), modelling can be a valuable tool for risk assessment of GMOs, acknowledging that modelling is complex even for presumably simple questions like the impact of Bt Toxins from transgenic maize (Dolezel et al., 2020; EFSA Panel on Plant Protection Products and their Residues, 2014; Fahse et al., 2018). While modelling ecological effects concerning gene drives is still in its infancy (Dhole et al., 2020), much research focuses on efficacy modelling. However, according to risk assessors, the view needs to be much broader than only efficacy.

The population-dynamic consequences of mate-choice, mating systems, and mating structure on gene drives are crucial in predicting the transgenic constructs’ probability and time to fixation and the release threshold for invading wild populations. Here the effects of mate-choice and mating systems are studied using deterministic ordinary differential equations. In contrast, the spatial mating structure uses a network model. Although we use different modelling frameworks for different mating complexities, the underlying gene drive model extends from a standard population genetic perspective. Gene drive systems have previously been categorized based on standard terminology; distortion in gamete proportions, fertility selection and viability selection (Verma et al., 2021). The need for such a common language across gene drive literature has been extolled by experts and regulators alike (Alphey et al., 2020). Here, we extend this approach by adding a generalizable understanding of the effect of some aspects of mating complexity on gene drive dynamics.

## Model and Results

As is typical for a functioning gene drive, we assume a diploid organism whose life cycle consists of three stages: zygote, adults and gametes. An adult produces gametes that combine to form a zygote. The zygote grows up to become an adult, and the cycle continues. We also assume that the organisms are diploid with two alleles for the gene of interest, the wild type allele (W) and the modified allele aimed to be driven (D). Hence, an individual can be either of the three genotypes: WW, DD and WD. Previous work has shown that the gene drive can arise if a drive carrying genotype undergoes distortion, viability or fertility selection that acts during the different life stages of an organism (Verma et al., 2021). Hence, one can categorize various gene drive systems based on pre-existing standard population-genetic terminology (distortion, fertility selection and viability selection). Manipulating the strength of these forces via the engineered construct influences the probability of inheritance, giving rise to gene drive (Verma et al., 2021). Gene drives can also be classified into two types based on the purpose of the release: modification and suppression drives. Suppression drives are aimed to reduce or completely eradicate the wild population, while modification drives are intended to replace the wild population with organisms carrying the gene drives. This article focuses on modification drives resulting from distortion and viability selection.

At the gamete level, distortion in gamete ratios can favour the transmission of the drive allele in the heterozygote. Gametes combine to form zygotes, but specific genotypes may become non-viable, distorting the expected Mendelian proportions of the zygotes. It can give rise to meiotic drive (Lindholm et al., 2016; Sandler and Novitski, 1957) and CRISPR based homing endonuclease gene drive (Noble et al., 2018, 2017). The engineered constructs that work principally by manipulating viability selection are those using zygotic toxin-antidote mechanisms as Medea (Beeman et al., 1992; Gokhale et al., 2014; Ward et al., 2011), Inverse Medea (Marshall and Hay, 2011) and Semele (Marshall et al., 2011). The general properties of engineered drives classified as such using standard population genetic terminology provide a clean comparative method when evaluating drives for risk assessment and deployment potential (Verma et al., 2021). Here, we subject the target population to three additional factors relevant to field populations, matechoice, mating structure and mating systems, to understand their effect on gene drive population dynamics (figure 1).

**Figure 1:**
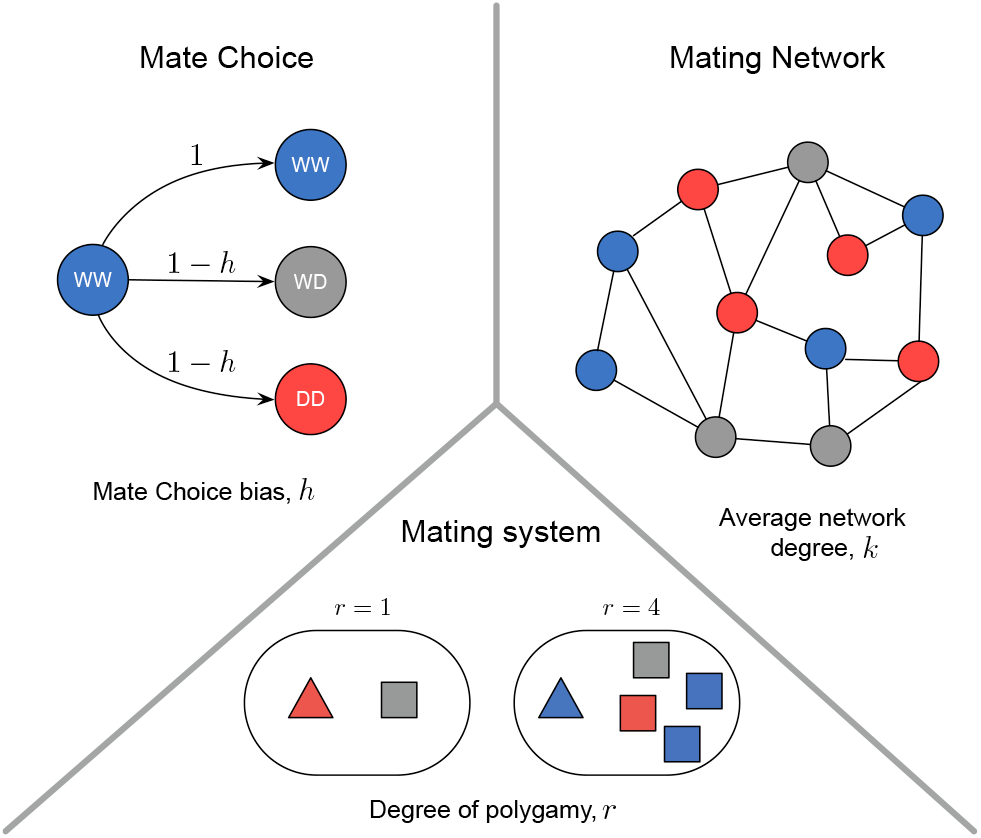
Pictorial representation of the three mating complexities: mate-choice, mating network, and mating system that can affect gene drive’s population dynamics. Blue, gray and red colours represent individuals with genotype WW, WD and DD, respectively. When there is no distinction between the two sexes, individuals are represented by circles, while triangles and squares denote individuals belonging to different sexes. Under mate-choice bias, the wild-type genotype (WW) are less likely to mate with drive carrying genotype (DD and WD). Mate-choice bias is denoted by *h* in our model, where (1 − *h*) is the mating rate between the wild-types (WW) and the transgenics (WD or DD). In structured mating, individuals mate and reproduce with other receptive individuals in their vicinity, and their likely interactions are modelled on a mating network of average degree *k*. The consequence of mating with one (monogamy *r* = 1) or multiple mating partners (polygamy, *r* > 1) on the gene drive dynamics is studied under the mating systems.

### Mate-choice

We first consider the null case where there is no gene drive and understand how mate-choice bias of wild-type against transgenic will affect the population dynamics. The mating rate among the wild-types is set to one. Similarly, the mating rate among the drive types is also one. Mate choice bias in our model is captured by the parameter *h* (figure 1). The mating rate among the wild-types (WW) and the drive types (WD or DD) is (1 − *h*). If *h* = 0, the wild-type (WW) are equally likely to mate with the drive carrying genotype (WD and DD). While if *h* = 1, the wild-type (WW) and the drive type (WD or DD) do not mate at all. During the exploration of parameter space (*h*), we work under the assumption that the wild-type genotypes are less likely to mate with individuals carrying the drive allele (WD and DD); therefore, 0 ≤ *h* ≤ 1. Our investigation of mate choice is motivated by the theoretical expectation that if variants of ornaments or behaviours that mate choice could act upon were linked to the same autosome where the gene drive is located and the cost of the drive is not zero (*c* ≠ 0) then mate choice could potentially evolve (Manser, 2015). It should be noted that despite substantial speculation and experimentation with autosomal natural gene drives, particularly the t-haploytpe in mice, there is no reliable evidence that mate choice has evolved (Sutter and Lindholm, 2016). However, behavioural traits that do appear to be linked to a drive locus have been recently described (Runge and Lindholm, 2021). Regardless, it is still potentially valuable to examine the relative sensitivity of various types of gene drive to the impact of mate choice should it evolve. This is not only for perspectives relating to the applied uses of gene drive, but also for the origin and evolution of natural drives elements. For simplicity, in our model, both sexes (male and female) of WW have an equal bias against mating with WD or DD. The rate of the production for the three genotypes assuming an infinitely large population and random segregation of alleles during meiosis is given by,

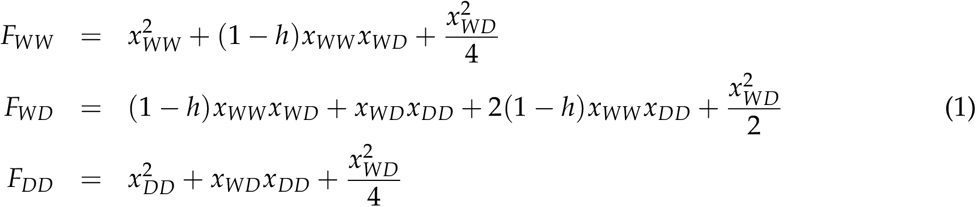

where *x*_*α*_ and *F*_*α*_ are the frequency and rate of genotype production respectively, and *α* ∈ (*WW, WD, DD*). The following set of differential equations governs the population dynamics of the genotypes in continuous time:

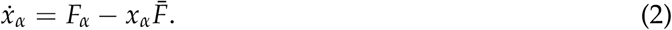

where 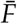 is the average fitness of the three genotypes:

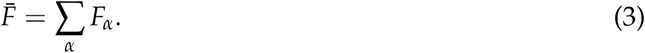

The frequencies of all genotypes is normalised to one.

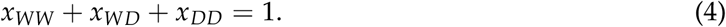

The above constraints on frequencies allow us to represent the dynamics of equation (2) on a de Finetti diagram. The frequency of the three genotypes (WW, WD and DD) without mate-choice (*h* = 0) converge to Hardy Weinberg equilibrium (Gokhale et al., 2014; Verma et al., 2021). When we introduce the mate-choice parameter into the rate equations (1), the dynamics deviate from Hardy Weinberg equilibrium and is governed by the fixed points that appear in the interior of the de Finetti diagram. In this context, a fixed point is a specific composition of the population 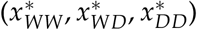 where the proportion of all the genotypes does not change. Specifically, where 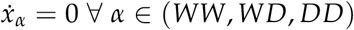. Primarily, there are two types of fixed points: stable and unstable. If the population is at the stable fixed point, a slight change in the population composition will bring the population to the stable fixed point. While in unstable fixed points, a small change will diverge the population composition away from an unstable fixed point. The position of these fixed points governs the overall population dynamics of a specific case. For example, population dynamics for a particular case of *h* = 0.9 is shown in the inset of figure 2A. The position of an unstable interior fixed point decides the evolutionary fate of the population.

**Figure 2:**
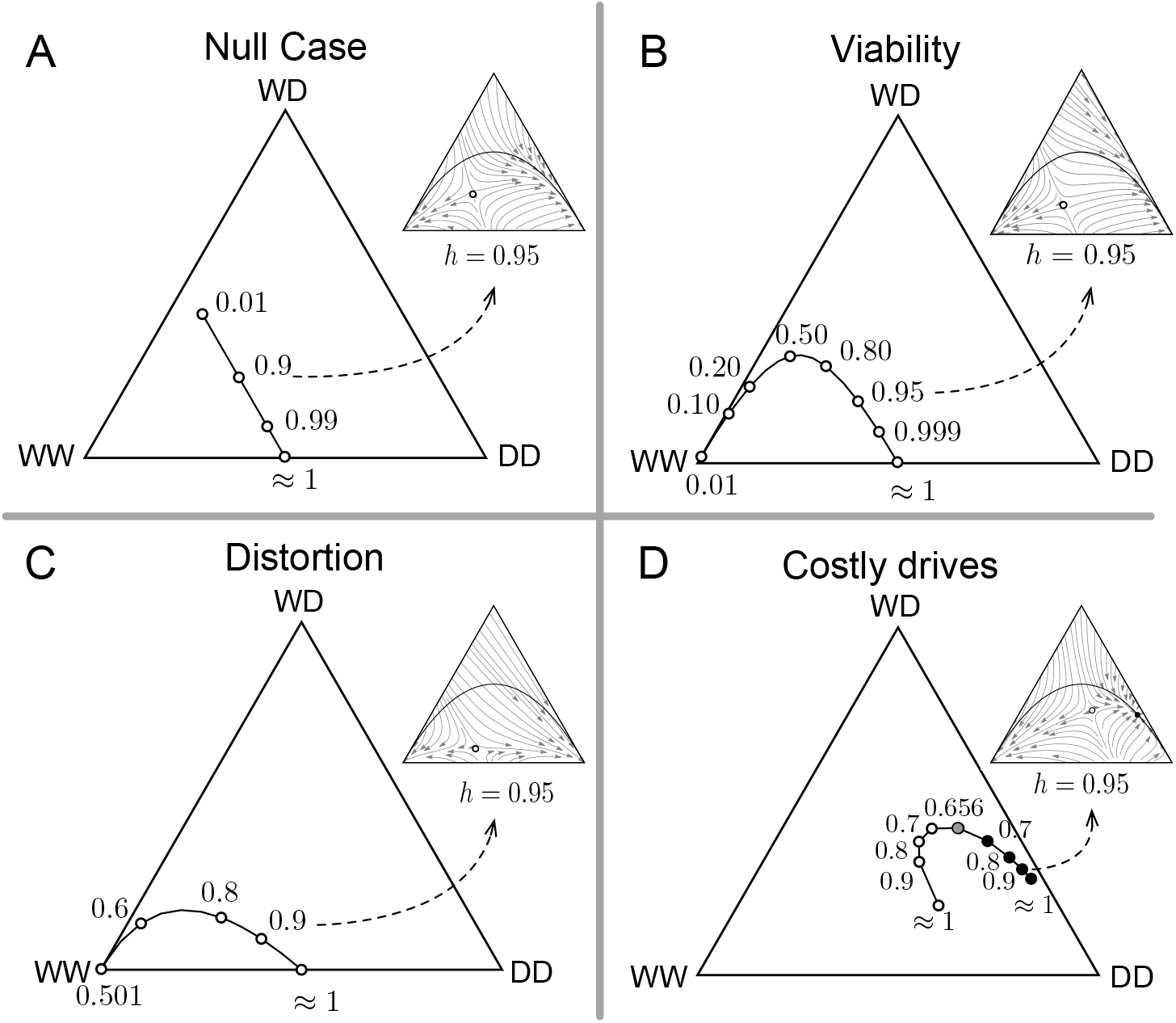
Effect of mate-choice bias (*h*) on the internal fixed point of the population dynamics without (null case) and with gene drives system based on viability selection (Medea), distortion and when they are costly. Fixed points appear in the interior of the de Finetti diagrams when the fitnesses of all the genotypes are the same. Open circles denote unstable fixed points of the dynamics, while closed black circles denote stable fixed points. The gray circle denotes the bifurcation point where both unstable and stable points emerge. The position of these fixed points changes with mate-choice bias (*h*) and hence the overall population dynamics, including the release threshold. Solid black lines show the trajectory of these fixed points for varying mate-choice parameter *h*. **(A)** Null case (without drive) considers the effect of matechoice alone on the population dynamics. **(B)** Medea drive efficiency is set to 100%, *d* = 1.0 **(C)** Distortion based drive is assumed to be fully efficient (probability *p* = 1.0) **(D)** Fitness cost of the drive allele is set to *c* = 0.2 and thus under a multiplicative cost model *f*_*WD*_ = (1 − *c*) and *f*_*DD*_ = (1 − *c*)^2^ with *f*_*WW*_ = 1. When other parameters are not changed their values are: *d* = 0, *p* = 0.5, *f*_*WW*_ = 1, *f*_*WD*_ = 1, *f*_*DD*_ = 1. The population dynamics for a specific case of *h* are shown in the insets for all panels.

In figure 2, we plot the positions and trajectories of these interior fixed points for different mate-choice (*h*) values under scenarios such as null case, viability selection, distortion, fertility selection. The null case is when only the effect of mate-choice is considered without any gene drive arising from viability selection, distortion, and fertility selection (figure 2A). Even under slight mate-choice bias (*h* = 0.01), the dynamics quickly deviates from the Hardy Weinberg equilibrium. An unstable fixed point (saddle point) appears in the interior of the de Finetti diagram. The threshold frequency of transgenic genotype (DD or WD) required for population transformation is closely related to the position of these unstable fixed points. The area to the left of the unstable fixed point is the basin of attraction of wild-type genotype. The trajectories of the initial conditions in this area lead to the extinction of the modified allele. In contrast, the area on the right is the basin of attraction of drive homozygotes (DD), leading to population transformation. Increasing the mate-choice bias (or as *h* increases from 0.01 to approximately 1), the position of the interior fixed point moves towards the middle of WW and DD line (figure 2A).

It implies that when the mate-choice bias increases, the threshold amount of transgenics (DD and WD) required to transform the wild-type population increases even without the gene drive.

### Mate-choice with Viability Selection (Medea)

Many toxin-antidote gene drive designs, including Medea, Inverse Medea, Semele, and designed under-dominance drive, exhibit viability selection (Beeman et al., 1992; Marshall and Hay, 2011; Marshall et al., 2011). In such systems, specific offsprings become non-viable during the zygote stage of the life cycle. We have focused on the Medea gene drive system in our analysis, where *d* measures the drive efficiency. In Medea gene drive, wild-type homozygous offspring of heterozygous mothers become non-viable (Akbari et al., 2014; Buchman et al., 2018; Gokhale et al., 2014; Ward et al., 2011). The rate of production of genotypes in the for Medea gene drive with the incorporation of mate-choice bias can be written as:

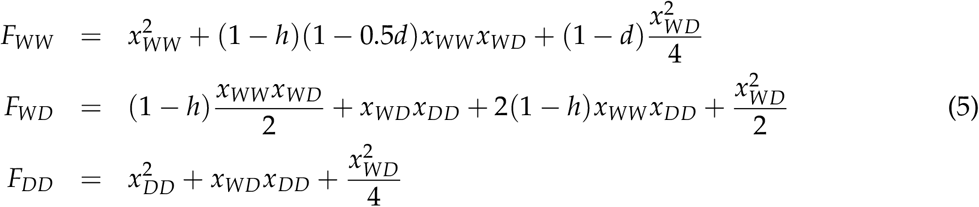

Figure 2B shows the position and trajectory of the unstable fixed point for viability selection based Medea gene drive with 100% efficiency, i.e. *d* = 1. The population dynamics equation can been derived using equation (2) and (5). When the mating rate between transgenic and wild-type decreases via *h*, the unstable fixed point moves towards DD vertex in the de Finetti diagram following a projectile trajectory (figure 2B). Hence here, mate-choice bias increases the threshold release of transgenics. For *h* ≈ 1, the number of transgenics released needs to be almost half the target population size for achieving total population replacement. These results are also consistent with the invasion condition of equation (A3) derived in appendix A for Medea gene drive.

### Mate-choice with Distortion

Here we will consider the case of distorted allele transmission in addition to mate-choice bias introduced by *h*. There are several distortion based gene drives, but here we will focus on a meiotic drive where the distortion efficiency is *p*. More specifically, *p* is the probability of transmission of drive allele from heterozygous parent to offspring. If *p* = 1, the gene drive system mimics CRISPR/Cas-9 based homing endonuclease drive with 100% efficiency (Noble et al., 2017). If a drive allele is transmitted from heterozygous parents with probability *p*, the rate of genotype production then changes to,

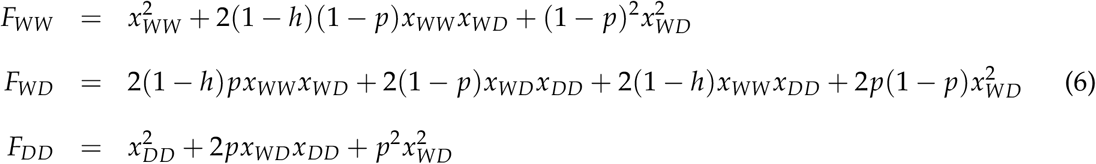

Again the population dynamics for the distorted case is given by equation (2), but the effective genotype production rate changes according to equation (6). In figure 2C we focus on the scenario when the distortion based gene drive such as meiotic drive or CRISPR drive with 100% efficiency (refer equation (6) for *p* = 1). We observe that the interior unstable fixed point only appears after the mate-choice bias becomes greater than 50% or *h* > 0.5, unlike viability based gene drive Medea (figure 2B & C). For *h* < 0.5, a small transgenic release is enough for population transformation to drive homozygotes (DD). Hence, the distortion based gene drives appear to be more robust against the mate-choice than viability-based gene drive Medea. These results are also consistent with the condition of invasion derived in appendix A for the distortion based gene drive (see equation (A6)).

### Costly drives

The relative number of offspring produced may differ because of the variation in the fertility of the mating pairs resulting from their genotypes especially if the drive confers a cost. The fitness component due to differential fitnesses is included in the parameters *f*_*α*_ where *α* ∈ (WW, WD, DD). The rate of the offspring production for the three genotypes is then given by,

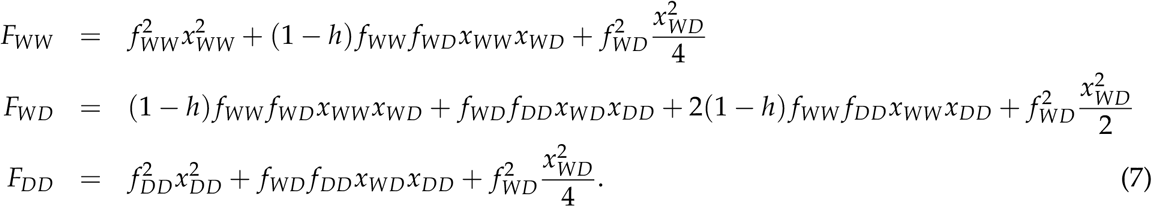

To observe the effect of fitness cost, we consider a scenario where *f*_*WW*_ = 1, *f*_*WD*_ = (1 − *c*), *f*_*DD*_ = (1 − *c*)^2^ for the dynamical equations derived using equation (7). Here, we assume multiplicative fitness cost where *c* denotes the fitness cost of the drive allele. The two internal fixed points appear only after substantial mate-choice bias *h* ≈ 0.656 (figure 2D). One of the fixed points is unstable, and the other is stable. Therefore, with the multiplicative fitness cost of the transgenic organism, due to drive-allele payload, mate-choice can result in the coexistence of all three genotypes. When *h* < 0.656, the global stable fixed point lies at the vertex of the wild-type population (WW); hence no amount of drive release can replace the wild population; however, complete fixation may not be a necessary aim in all applied scenarios.

Besides understanding the impact of mate-choice on the population dynamics, we also indirectly probe the threshold release fraction of transgenic organisms required for complete population replacement relative to the target population size. In figure 3, we numerically calculate the threshold frequency of drive homozygotes (DD) necessary to invade a population consisting of wild-types (WW). We evaluate the impact of mate-choice bias (*h*), gene drive efficiency and fitness cost for two gene drive systems, namely meiotic drive and Medea. Figure 3A shows that the mate-choice bias increases the invasion threshold frequency of DD required for complete population replacement for Medea drive. The threshold frequency of DD also slightly increases with decreasing drive efficiency. The change in threshold frequency due to drive-efficiency reduces for increasing bias in mate-choice. The release threshold is close to zero for lower mate-choice bias, represented by the heatmap’s light colour. The position of fixed point for the case of 100% drive efficiency (*p* = 1 and *d* = 1) for both figure 3A & B corresponds to the scenario studied in 2B & C respectively. Lower mate-choice and sufficiently high distortion probability do not change the threshold frequency for the distortion-based drive. The region in the heatmap where a minimal transgenic release can transform the population is significantly high for the distortion-based drive than Medea drive. When the mate-choice bias is high enough (*h* > 0.5), an increase in distortion probability only slightly decreases the invasion threshold of DD. In this regime (*h* > 0.5), a substantial frequency of DD is required for the wild-type population to be invaded even for a very high distortion probability.

**Figure 3:**
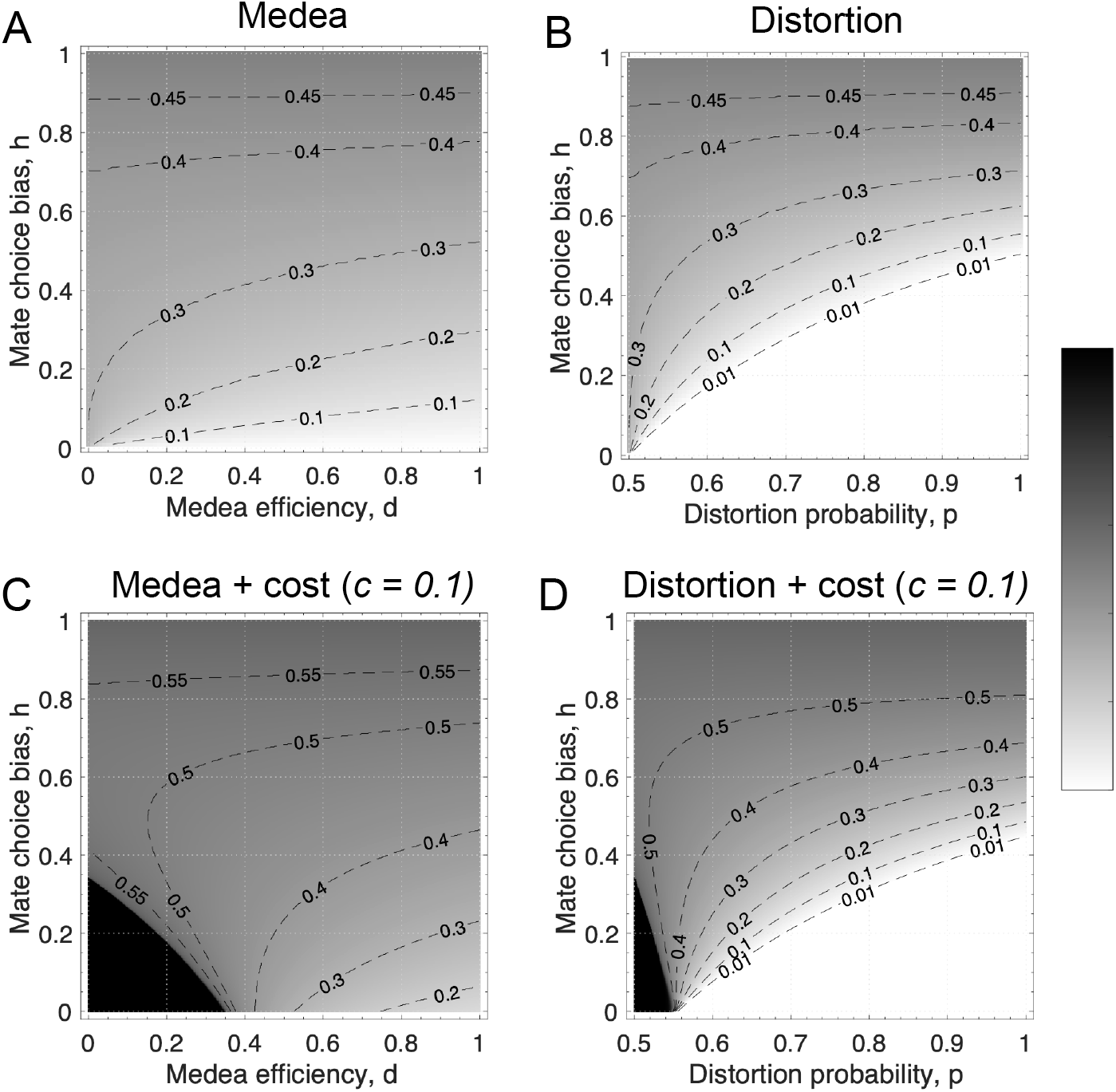
Heat-map shows the threshold frequency of drive homozygotes (DD) required to invade a population of wild-type homozygotes (WW) with respect to variation in mate-choice bias (*h*) for the following gene drive systems: Medea and distortion based drive. Black dashed lines correspond to the contour lines showing the threshold frequency of drive homozygotes (DD). (**A**) Medea gene drive with no fitness cost i.e. *c* = 0. (**B**) Distortion based gene drive with no fitness cost to drive i.e. *c* = 0. (**C**) Medea gene drive where the fitness cost due to drive allele is *c* = 0.1 hence *f*_*WD*_ = 0.9 and *f*_*DD*_ = 0.81. (**D**) Distortion based gene drive where the fitness cost due to drive allele is *c* = 0.1 hence *f*_*WD*_ = 0.9 and *f*_*DD*_ = 0.81.

In figure 3C & D corresponds to the case when there is a fitness cost for the drive carrying organism (*c* = 0.1 hence *f*_*WD*_ = 0.9 and *f*_*DD*_ = 0.81). Fitness cost leads to an increase in the invasion threshold frequency for both the gene drive systems overall. Moreover, any DD release is insufficient to invade the wild-type population for inefficient drives under low mate-choice bias. The dark colour represents this region in the heatmap. Interestingly, increasing the matechoice bias can facilitate the invasion by DD even for less efficient drives. The distortion based gene drive appears to be more robust against the ecological stress of mate-choice bias even when considering the fitness costs.

### Mating systems

Gene drive technology relies on sexual reproduction between the mating pairs. Most of the target species of interest have a polygamous mating system instead of the commonly assumed monogamous mating system (Moro et al., 2018; Rode et al., 2019). As introduced in the previous section of mate-choice, the model is modified here to incorporate this aspect of the mating system. In this model, we will consider two separate populations of the two sexes. We assume that the offspring of both sexes are produced in equal proportion. The frequency of male and female genotypes are denoted using *x*_*i*_ and *y*_*j*_. There are three possible genotypes: wild-type (WW), drive heterozygotes (WD) and drive homozygotes (DD). Let us consider the mating system when one male mates with *r* females. Hence *r* = 1 represents the monogamous mating system while *r* > 1 corresponds to the polygynous mating system. The following set of equations gives the frequencies of the genotypes produced with the polygamous mating system, as the equation holds for both males and females (with a change in variable *x*_*i*_ and *y*_*j*_):

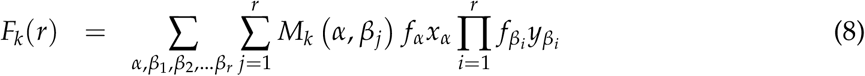

Here, *M*_*k*_ (*α, β*_*j*_) is the proportion of genotype *k* produced from the mating between a male of genotype *α* and a female of genotype *β*_*j*_. *α* and *β*_*j*_ are dummy indexes for any of the three genotypes WW, WD or DD. The elements of the matrix *M*_*k*_ (*α, β*_*j*_) will depend on the gene drive type as well. Matrix *M*_*k*_ for Medea (equation (A7)-(A9)) and distortion based gene drive system (equation (A10)-(A12)) is given in appendix A. The summation over *α* and *β*_*j*_ is carried out over the set of all genotypes (WW, WD, DD). We have also assumed a polygamous mating system of mating ratio *r*, i.e. one male mates with *r* female or vice-versa. Equation (8) may be interrupted in parts as selecting a male of genotype *α* and selecting *r* females of genotype *β*_1_, *β*_2_, …, *β*_*r*_. Finally, the contribution of all possible matings in producing genotype *k* is summed up.

Simplifying equation (8) by expansion formula for multinomial expression yields,

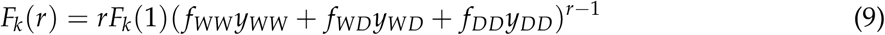

The following set of differential equations governs the population dynamics of the genotypes in continuous time:

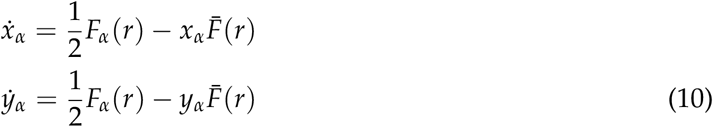

where 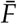 is the sum of rates of genotype production:

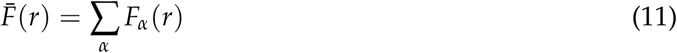

The total population of both males and females remains constant and sum up to unity.

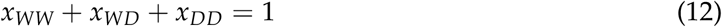

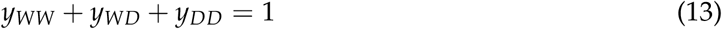

In equation (9), *F*_*k*_(*r* = 1) and *F*_*k*_(*r* > 1) is the production rate of genotype *k* for monogamous (*r* = 1) and polygamous (*r* > 1) mating system respectively. It implies that the equilibrium population dynamics for both monogamous (*r* = 1) and polygamous (*r* > 1) mating systems, even with gene drives, are equivalent. In other words, the final population composition of the genotypes remains the same for both polygamous and monogamous mating systems. Previous studies without any gene drive also support that the equilibrium dynamics for monogamy and polygamy remain the same (Karlin, 1978; O’Donald, 1980). However, the difference lies in the relative time to reach population equilibrium. It can be shown that after simplifying the equation (10) obtained for *r* > 1, the rate of increase of different genotypes is equivalent to the case of monogamy (*r* = 1) with rescaled time. The expectation is that the gene drive will spread faster in polygamous mating species compared to monogamy (Moro et al., 2018). Hence, the time required for the drive allele to spread through the population should increase for the monogamous mating system. Our result also supports the expected outcome. Here we quantify the same.

If we first look at the case where there is no fitness cost of the gene drive, and only the efficiency of the two gene drive system based on distortion and viability selection are varied. Figure 4 left and center panels show that gene drive will spread faster for species with a high degree of polygamy (r). In the online material provided we show that the distortion-based gene drive will spread faster than the viability-based drive. The time for the gene drive to reach 99% frequency is an order of magnitude higher for Medea drive compared to CRISPR homing drive or meiotic drive. A higher degree of polygamy (*r*) reduces the time required to reach critical drive frequency (99%) for both the gene drive system.

**Figure 4:**
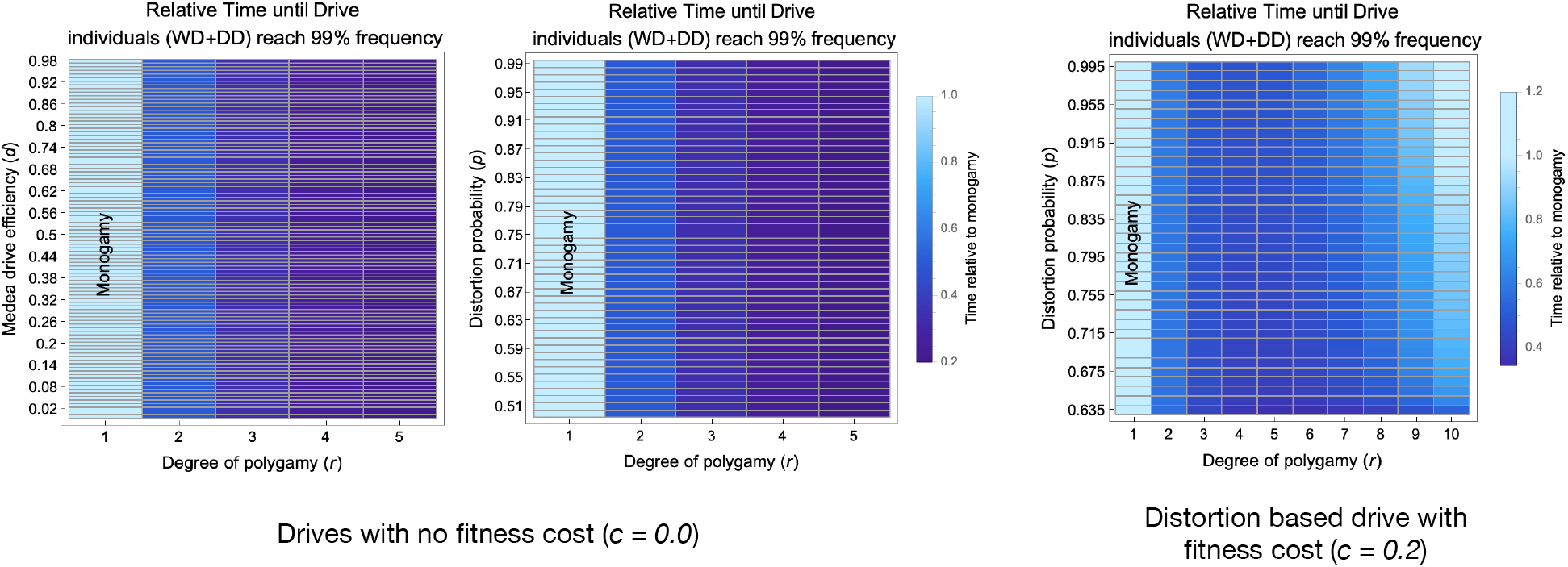
Effect of mating system and drive efficiency on the time for the drive allele to reach 99% frequency. We start from a population consisting of the 99% wild-types (WW) and 1% the drive heterozygotes (DD). The population is evolved until frequency of drive allele carrying individuals (WD + DD) reaches 99% (“fixation”). The left and centre panels do not include a fitness cost attributable to the drive. The left panel shows the fixation time for different degrees of polygamy across the range of Medea efficiency normalised by the time taken in the monogamy case. The central panel shows the fixation time for different degrees of polygamy across the range of distortion efficiency normalised by the time taken in the monogamy case. Both panels show a linear dependence on the degree of polygamy which continues for higher values of *r* (not shown - available in online code). When including a fitness cost towards carrying a drive allele *c* = 0.2, the fixation time shown a non linear dependence on the degree of polygamy. For low and high degrees of polygamy it takes longer to fix than for intermediate levels of polygamy. There is also a slight dependence on the efficiency of the distortion drive where lower probability of distortion is still faster for intermediate levels of polygamy as compared to the monogamous case.

Figure 4 panels for costless drives show that the relative time required for the drive allele to reach 99% frequency is rescaled exactly by a factor of 1/*r* for the polygamy relative to the monogamous mating system (e.g. time relative to monogamy is 0.2 for degree of polygamy *r* = 5). This is in line with the relation obtained in equation (9). When *f*_*WW*_ = *f*_*WD*_ = *f*_*DD*_, the production rate of offspring for polygamy is *r* times that for the monogamous mating system. But, when we have a fitness cost *c* for carrying a drive allele, the relation between the time to reach 99% frequency and degree of polygamy becomes more complex (figure 4). An increase in the degree of polygamy first decreases the relative time to reach the drive allele’s critical frequency (*r* = 2 and *r* = 4), but a further increase in the degree of polygamy (*r* = 6, 8, 10) elevates it. In figure 4, it can also be noted that when the distortion probability is low (*p* < 0.625), the costly drive allele is not able to invade the wild-type population. Thus in the right panel we explore values of distortion probability above this threshold. This is in congruence with the condition of invasion derived for the monogamous case in equation (A6) in Appendix A.

The above result can be understood from the equation (9) where the fitness cost makes the factor (*f*_*WW*_*y*_*WW*_ + *f*_*WD*_*y*_*WD*_ + *f*_*DD*_*y*_*DD*_)^*r*−1^ less than one. The factor (*f*_*WW*_*y*_*WW*_ + *f*_*WD*_*y*_*WD*_ + *f*_*DD*_*y*_*DD*_)^*r*−1^ decreases exponentially with increasing level of polygamy *r*. Hence the time is rescaled by the factor of 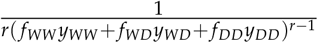 effectively. The time first decreases when dominated by 1/*r* with an increase in *r* but later on decreases when dominated by 1/(*f*_*WW*_*y*_*WW*_ + *f*_*WD*_*y*_*WD*_ + *f*_*DD*_*y*_*DD*_)^*r*−1^. When the fitness cost is *c* = 0.2, the relative time until the drive allele reaches 99% frequency with respect to monogamy decreases for *r* = 2 and *r* = 4, but then it starts to increase for *r* = 6. For *r* = 8 and *r* = 10 spread of gene-drive becomes slower compared to monogamy. Another way to understand the results is that the rate of production genotype DD first increases up to a point for increasing level of polygamy *r* but later decreases for moderate fitness cost (figure 5). Hence the time in spreading gene drive is lowest for intermediate levels of polygamy. Further increase in the degree of polygamy reduces the production of DD and therefore increases the time to spread the drive allele.

**Figure 5:**
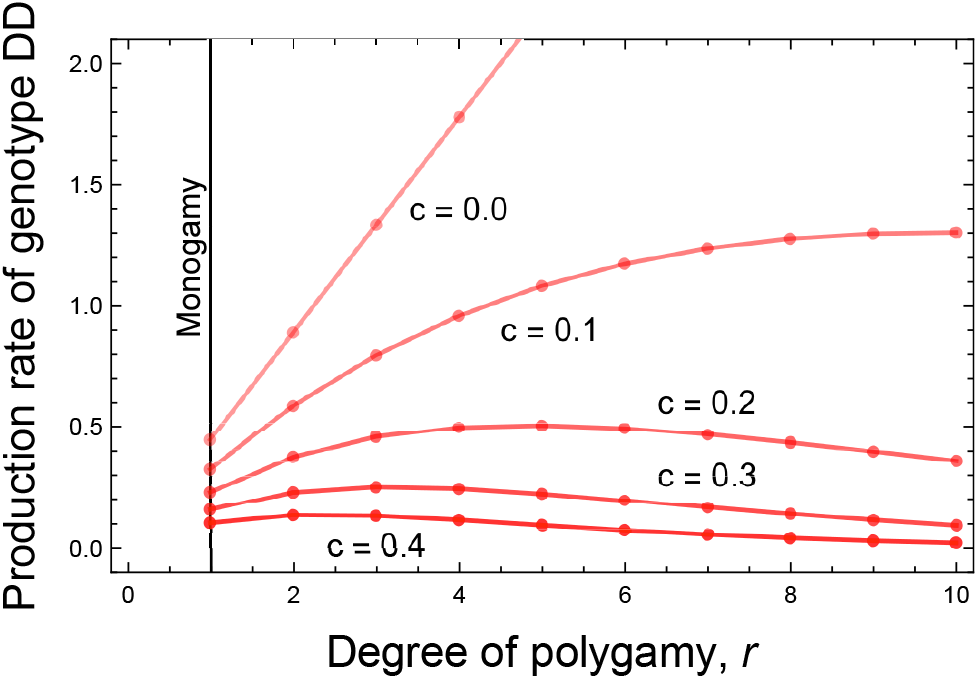
Effect on the rate of production of DD genotype with increases in degree of polygamy (*r*) for different fitness cost (*c*). We start from a population with an equal abundance of all three genotypes with 100% drive efficiency of distortion-based gene drive for different fitness costs. In essence, we plotted equation (9) for varying *r* and *c* keeping *x*_*WW*_ = 1/3, *x*_*WD*_ = 1/3, *x*_*DD*_ = 1/3 and *p* = 1. Increasing the fitness cost of the drive allele decreases the overall production of the DD genotype. For a moderate level of fitness cost, production of genotype DD first increases up to a point for species with a higher level of polygamy but then started to decrease.

### Spatial network interaction

The population dynamics of CRISPR based homing endonuclease gene drive have been extensively studied for well-mixed infinitely large (Noble et al., 2017) and finite populations (Noble et al., 2018). But most species occur in a partially heterogeneous landscape where they interact and mate with other individuals in their vicinity. Hence, a network-based population is an appropriate framework to model dynamics in such structured populations.

We considered a structured population of *n* individuals. The individuals live on a random network with an average degree of *k*; thus, each individual has *k* connections on average. Here *k* controls the number of mating opportunities and the level of competition for an individual. The population is updated via a death-birth process (figure 6) described as follows: First, an individual is chosen randomly for death. Then two parents are selected, who are neighbours of the dead individual with probability proportional to their fertility fitness. According to their genetic archetype, the selected parents contribute their gametes, where other genetic effects like distortion can come into play. The combination of these contributed gametes forms the offspring that replaces the dead individual in the network. The population is updated until it fixes to all WW or all DD states.

**Figure 6:**
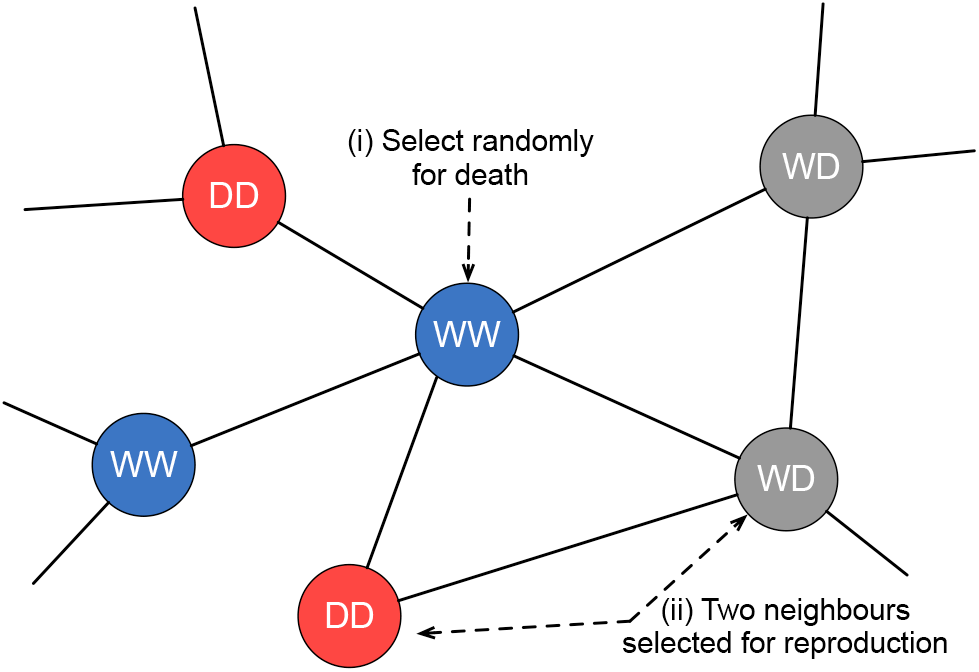
Spatial model explaining the population update mechanism. The blue, gray and red colours represent individuals of WW, WD and DD genotype, respectively. Population update happens in 2 steps: firstly, a random individual is selected for death. This step creates space at that particular network position. Secondly, two random neighbours of the dead individuals are chosen as parents to produce offspring. The genotype of the offspring is determined from the parents, and it replaces the dead individual.

In figure 7, we exhibit the stochastic network model by running the simulation several times and plotting fixation probability and conditional fixation time with variation in the average number of interacting individuals per site (represented by *k*). We also studied the impact of increasing the number of released transgenic (WD and DD) and different genotypes (WD and DD). Here, *k* controls the number of mating opportunities and competition during the birth process. When *k* increases, the fixation probability of DD decreases mainly due to higher competition during the birth update per site (figure 7A). As expected, distortion probability has a positive impact on the fixation probability of DD. The effect is more pronounced for lower values of an average degree since the heterogeneity in the number of connected individuals is also high for this case. Fixation probability also increases as the number of released DD increases (figure 7A). Releasing DD transgenics has a lower chance of getting fixed than a WD release (figure 7B). This is because the fitness cost of DD is relatively high compared to WD (*f*_*WD*_ = 0.50, *f*_*DD*_ = 0.25). If the fitness cost is negligible and the drive efficiency is high, the release of the DD genotype is expected to fix the gene drive with a higher probability. The effect on fixation probability by the release of WD compared to DD becomes more pronounced with the increase in average degree *k* (figure 7B). It increases first with an increase in the release number of transgenic, attains a maximum and decreases later. We present the fixation probability for a gene drive with a high fitness cost in part motivated by observations that many, but not all natural autosomal gene drives, e.g. *Mus musculus* t-haploytpe and the *Drosophila melanogaster* SD alleles are frequently homozygous male sterile. However the impact of the average degree of the network is indeed a result of the cost as it brings about a density dependent aspect to the number of released individuals. The effect is clarified in the supplementary figure A2 where we show the importance of discerning the fitness costs of a drive before releasing into a sparsely or densely connected networks.

**Figure 7:**
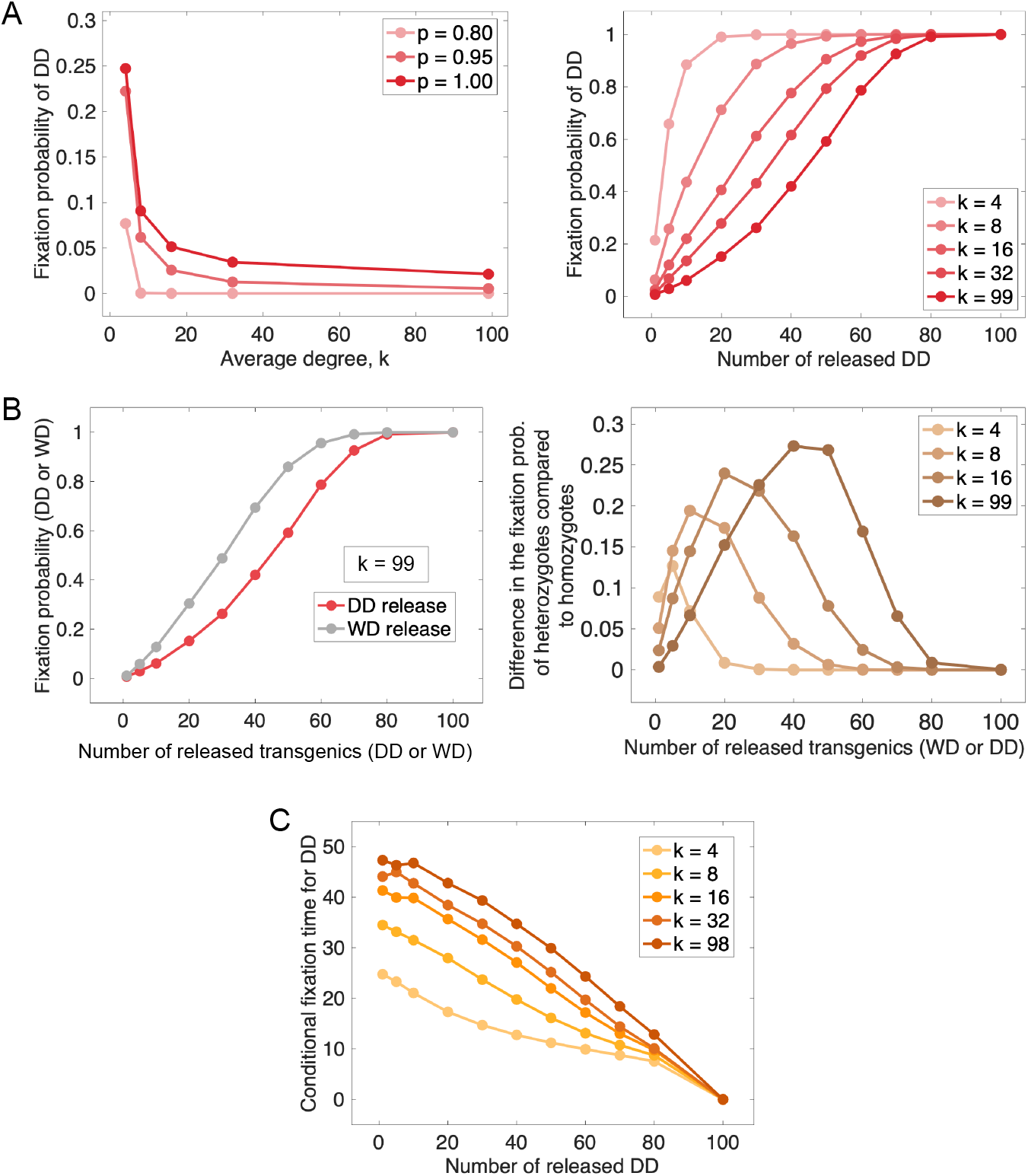
Fixation probability and conditional fixation time of DD with variation in average degree *k*, distortion probability *p* and initial number of released transgenic individuals WD or DD. **(A)** Plots show the fixation probability of drive homozygotes starting from a single DD individual against average degree *k* (left) and the number of released DD (right) for different values of *p* and *k* respectively. **(B)** Left: Fixation probability is plotted against the number of released DD and WD for a complete graph (*k* = 99). Right: the difference between the fixation probability of WD and DD release is plotted against the number of released transgenic for varying average degree *k*. **(C)** Shows the average number of generations when the drive individuals get fixed in the population against an initial number of released DD with varying average degree *k*. A generation consists of *n* death-birth step. Hence in a generation, the whole population is updated on an average. All simulations use a population size is *n* = 100 and 10, 000 trials to estimate fixation probability and conditional fixation time. The distortion probability and fitness cost are fixed to *p* = 0.95 (except panel A left) and *c* = 0.5, respectively. In all of our simulations for the release nodes of transgenic are chosen at random.

We also plotted conditional fixation times against the size of transgenic (DD) releases for various values of network degree (figure 7C). Unsurprisingly, this confirmed that the conditional fixation time is fastest for larger transgenic releases. Interestingly, conditional fixation time is slowest in populations with substantial structure (lower values of k). The broad range of values of conditional fixation times for the smaller release sizes reflects the increasing probability of stochastic loss of D alleles as population structuring increases.

## Discussion

Gene drive is one of the tools of synthetic biology that has the potential to transform whole wild populations. The transformation exploits and modulates one of the foundational tenets of evolution - the inheritance of traits through sexual reproduction. Thus, variation in the reproductive biology and the mating behaviour of the target species can affect the eventual spread of the gene drive. The analytical approach and terminology used in this study were selected to facilitate comparison between different drive mechanisms within a common comparable framework (Verma et al., 2021). While previous studies emphasised the evolution of resistance to gene drives (Price et al., 2020), we have examined some of the ecological assumptions related to mating systems and their effect on the potential outcomes of a drive release (Champer et al., 2018; Noble et al., 2017; Unckless et al., 2017). These include factors related to life histories and social interactions relevant under field conditions, namely mate-choice, multiple matings, and spatial aspects of the mating network. Our work focuses on single locus gene drives located on autosomes in diploid organisms. From an applied perspective aimed at using gene drives to manipulate populations, this equates to modification drives (not suppression drives that seek to reduce population sizes). We describe circumstances where the explored factors can substantially influence predictions for the release thresholds of transgenics required for a successful invasion and the possible fixation times.

First, we considered a monogamous situation that deviates from panmixis as individuals actively choose partners based on the presence or absence of a transgenic drive allele D (mate choice). For any drive linked to ornaments or behaviours that could form the basis of heritable mate choice, a mate-choice bias can be envisioned with a significant effect on the release threshold of a gene drive, as shown in figure 3. Inefficient drive and fitness costs due to drive-payload aggravate the situation. The predicted threshold release is drastically different from a situation with no mate-choice bias figure 2. This finding is important to estimate the drive efficacy and is highly relevant for the risk assessment of the drive. Comparing different drive approaches, we found that distortion based gene drives fare much better than drives based on viability selection under the potentially more ecologically realistic conditions of mate choice. While the regulatory importance of these insights may be limited due to the often unknowable capacity for mate choice to evolve, they may inform what parameters might be helpful to monitor subsequent to any release.

Next, we considered the potential for multiple matings for both females and males. Under the explored scenarios, the final evolutionary outcome of the spread of the gene drive (distortion or Medea drive) for a polygamous mating was the same in our simulations as that of the monogamous system. Even the species with a higher degree of polygamy will converge to the same evolutionary fate for a given gene drive system. However, the time needed for the spread of the drive gene will be affected by the level of multiple matings and the fitness cost linked to the drive. Time to fixation will be smaller for a higher degree of polygamy without any fitness cost (figure 4). However, a moderate fitness cost under different polygamy levels will trigger a nonlinear outcome for the time till drive establishment. This non-linearity is because the production rate of drive homozygote first increases but later decreases in line with the degree of polygamy for moderate fitness cost (figure 5). Hence, the drive gene is expected to spread faster for species with intermediate levels of polygamy when there is an associated fitness cost of the drive allele. Regarding mice, which are one of the focus organisms of gene drive applications, these primarily utilise suppression drives that are not considered here. It is, however, still interesting to note that the value of *r* = 2 that is applied for island mice (Birand et al., 2022), would imply a near 2 fold difference in fixation time compared to situations of monogamy, which may be significant in circumstances where alternative non-genetic approaches may be available to achieve the same goals.

Last, we examined the spatial implications of finding mating partners (mating opportunities) on the model outcome. To this end, the framework developed for the spatial mating interaction can be applied to any diploid population, regardless of the presence of a gene drive. Considering a finite population on a network allows us to understand the probable outcomes of gene drive release. A finite population leads to stochastic fluctuations in the frequencies of the genotypes resulting in different outcomes for the same initial conditions. We found that the spread of transgenic release is lowered when individuals, on average, have more mating opportunities and intra-sexual competition. Thus the fixation time for the transgenic increases with an increase in the average degree of the mating network. Concerning the question of how the connectivity of mating network varies in wild populations, it is reported that selective pressures under which species evolve shape their network structure in the environment (Pinter-Wollmann et al., 2014). Hence, changes in environmental conditions such as resource availability, seasonal effects, selective pressure, and life-history traits can impact the network structure. Within a species, variation at the individual level can also lead to heterogeneous connectivity. Species with sparsely connected individuals on the mating network have a higher chance of fixing drive genes in a shorter time. We also observe that the success in fixation of drive homozygotes can be mitigated by releasing more transgenic individuals. Furthermore, when the fitness cost associated with carrying a drive allele payload is high, releasing drive heterozygotes instead of homozygotes would result in a higher chance of gene drive fixation (figure 7B). Our model of network dynamics uses a death-birth process. Alternatively it is also possible that we first select a connected mating pair for birth that replaces an individual connected to either parent for death. In models of asexual reproduction on networks, such birth-death vs death-birth models can show qualitative differences in fixation properties (Hindersin and Traulsen, 2015; Kaveh et al., 2015; Zukewich et al., 2013). Exploring such alternative processes would require a thorough understanding of the network dynamics of the target species.

While our findings can collectively be viewed solely in applied uses of synthetically engineered drives, they may also inform the origin and evolution of natural gene drives in animals. Despite the intense focus on synthetic gene drive and analogies with described gene drive elements from nature, three natural elements fit the modelling described here. They are: autosomal in a diploid organism, single copy, do not alter population sex-ratios or involve genomic transposition. The three natural drive elements are described in Table 1. In contemporary wild populations, these drives can be detected as polymorphic (with a frequency < 1). None are associated with mate choice. With the strong caveat that three is a minimal sample size, there is an absence of information on how many drive elements may have evolved and risen to fixation in species (rendering them effectively undetectable except by inter-specific crossing). It is still interesting to speculate if the findings of this study provide any insight into the origin of natural gene drives and the probability that when they arose in the first individual, they spread or not. Our study would indicate that the greatest probability of a new gene drive allele increasing from a low initial frequency would be in species with low degrees of polygamy and a highly structured population (low k). This would particularly be the case for the *Drosophila melanogaster* SD alleles and mouse t-haplotypes where the cost of the drive (*c*) appears to be high. To gain confidence in these statements, it will be necessary to estimate the relative frequency of gene drive elements in a range of animals with different ecological properties. However, given the historical difficulty in detecting autosomal natural gene drive elements, it may take some time, even with the drop in genotyping costs, for extensive drive studies to take place outside intensively studied model organisms.

**Table 1:** Examples of natural drive elements with properties simulated in this study.

| Species | Drive element | Drive type & parameters | Ecological properties |
| --- | --- | --- | --- |
| *Tribolium castaneum* | Medea | Viability drive $d$ | Probably low $k$, highly polyandrous<br>(Pai and Yan, 2020) |
| *Drosophila melanogaster* | Selfish Segregation Distorter (SD) | Distortion drive $p, c$ | Probably low $k$, Polyandry with some re-mating latency (Singh and Singh, 2004) |
| *Mus musculus* | t-haplotype | Distortion drive $p, c$ | Probably low $k$, Polyandry ($r = 2$, (Birand et al., 2022)) |

In this study, we have decided to focus on three factors related to the mating complexities of the target species. Still, many other ecological and environmental factors can impact the spread of gene drives. Known factors include the age structure of a population, spatial landscape and seasonality (Eckhoff et al., 2017; Huang et al., 2009, 2011; North et al., 2013, 2019, 2020). Gene drive behaviour and the interactions of a drive released in a complex ecosystem over long time periods are highly complex. Navigating this ecological complexity may seem insurmountable (Levin, 2003). However, we will face a similar control problem for any technology aiming to intervene in complex systems. As such, it is not workable to address *in silico* all possible ecological and evolutionary pressures and scenarios that an engineered system will meet in the real world (Denton and Gokhale, 2019; Lindvall and Molin, 2020). Undoubtedly, modelling will play a key role in understanding drive spread. Our study emphasises that modelling needs a more ecological reality to predict the drive behaviour. Identifying and collecting necessary information on the effect of primary ecological and evolutionary pressures will be thus crucial to access the risk before any field deployment (James et al., 2018; Long et al., 2020; of Sciences Engineering and Medicine, 2016).

## Conclusion

Most of the gene drive modelling exercises focus on drive spread under simplified conditions such as panmixis. In this study, we tested whether more complex assumptions, characteristic of many species and mating systems, justify the use of simplified assumptions. The results show that ecological factors related to mating can substantially change drive spread and, as supposedly many other ecological factors, may strongly impact gene drive systems’ temporal and spatial dynamics. Modelling may be used to predict gene drive spread and thus to assist in risk assessment. In this case, mating-related parameters, as all critical assumptions related to the ecology of the species, need to undergo a reality check. In a wider sense, the new modelling framework, including tools for analysing spatial interactions or multiple matings, are generic and have the potential to be applied to any diploid population, independent of gene drive applications.

## Acknowledgments

The work has been supported by funds from the Max Planck Society and is part of the R &D project “Risk assessment of synthetic gene-drive applications” (FKZ 3518 84 0500) funded by the German Federal Ministry for the Environment, Nature Conservation and Nuclear Safety commissioned by the Federal Agency for Nature Conservation (BfN), Germany.

## Data and Code Availability

All data and simulation codes for generating figures are available in an online repository (https://anonymous.4open.science/r/genedrives_mating-6841/).

## Appendix A: Additional Methods

### Invasion condition for Medea drive with Mate choice (h)

If we consider the case of Medea gene drive with fertility selection. The rate of production of the three genotype is given by the combination of equation (5) and (7),

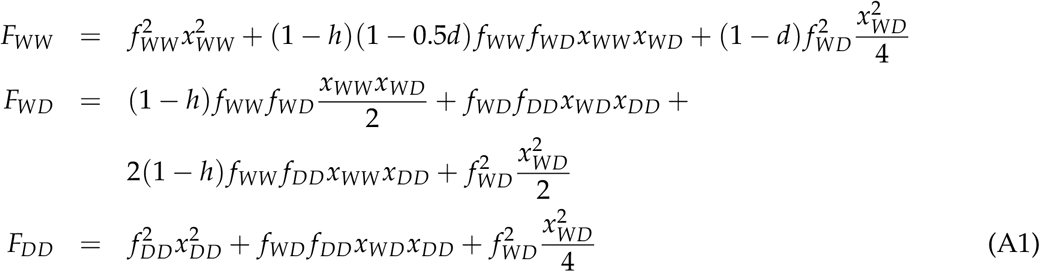

The rate of change of frequencies of each genotype is still given by equation (2). We use the constraint on the frequencies of the three genotypes in equation (4) to reduce the population dynamics of the genotypes to two variables after replacing *x*_*WD*_ = 1 − *x*_*WW*_ − *x*_*DD*_ in equation (2). The drive will not invade the wild-type population if both the eigenvalues of the dynamical system are negative. Eigenvalues can be deduced from the Jacobian matrix (*J*_*d*_) of the system at (*x*_*WW*_, *x*_*WD*_, *x*_*DD*_) = (1, 0, 0),

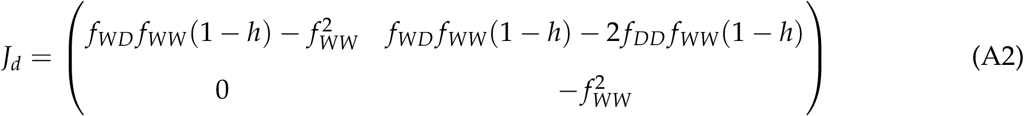

Hence, Medea gene drive can invade a population of wild-type if

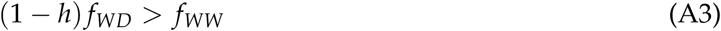

Note that the above invasion condition is independent of the efficiency of the Medea gene drive (*d*). This condition is plotted in figure A1 (left panel).

**Figure A1:**
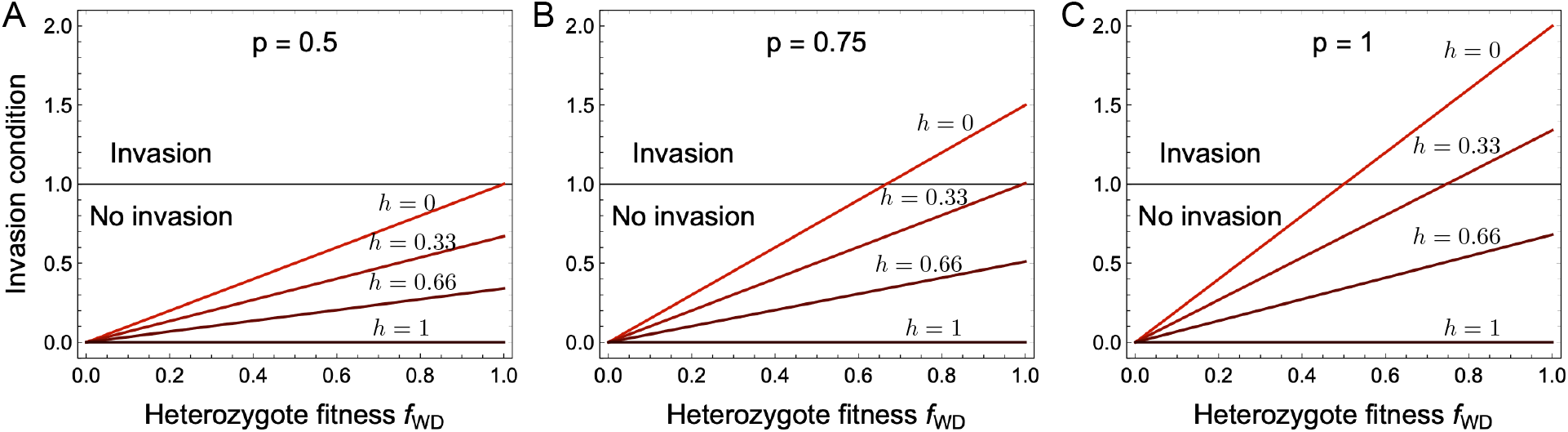
Invasion condition with varying mate-choice bias (*h*) against heterozygotes fitness. *f*_*WD*_. Here, invasion condition refers to the ability of the drive to invade a wild-type population from low frequency, with an invasion condition of greater than 1 being necessary for success. (**A**) Medea drive or no distortion, *p* = 0.5. Wild-type population cannot be invaded for any value of mate choice bias, *h*. (**B**) Distortion based gene drive with *p* = 0.75. Wild-type population can be invaded if there is no-mate choice bias *h* = 0 and *f*_*WD*_ > 2/3. (**C**) Distortion based gene drive with *p* = 1. Wild-type population can be invaded if mate choice bias not very high i.e. for *h* = 0 and *h* = 0.33.

### Invasion condition for Distortion drive with Mate choice (h)

Consider the scenario of distortion based gene drive with fertility selection. The rate of production of the three genotypes will then be governed by the combination of equation (6) and (7),

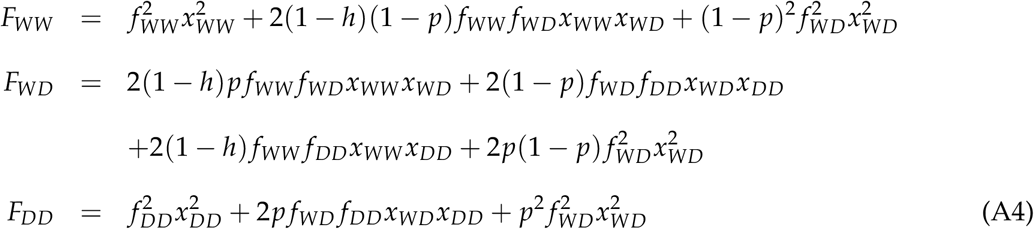

Similar to the Medea gene drive scenario, the population dynamics of the above system can be written in the form of two variables *x*_*WW*_ and *x*_*DD*_ using equation (4). The Jacobian matrix (*J*_*m*_) of the system at (*x*_*WW*_, *x*_*WD*_, *x*_*DD*_) = (1, 0, 0) is given by

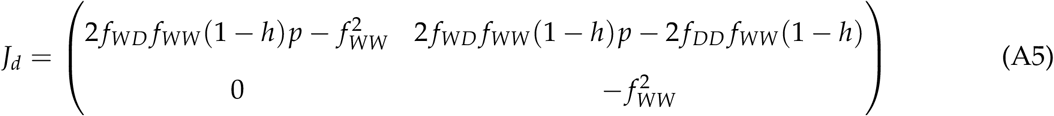

From the condition on the eigenvalues, the gene drive can invade wild-type population if,

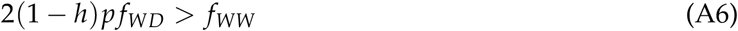

as shown in figure A1 (centre and right panels). Note that when there is no mate choice (*h* = 0) the above condition reduces to the invasion condition derived by Noble et al. (2017) for CRISPR gene drive.

### M_k_ (α, β_j_) in equation (8) for Medea and Distortion Based Gene Drive

#### Medea Gene Drive

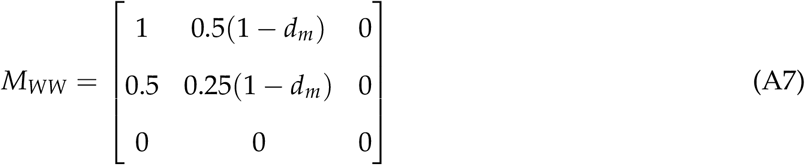

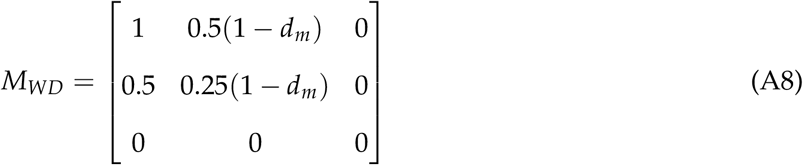

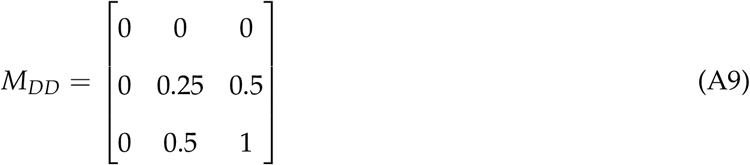

#### Distortion Based Gene Drive

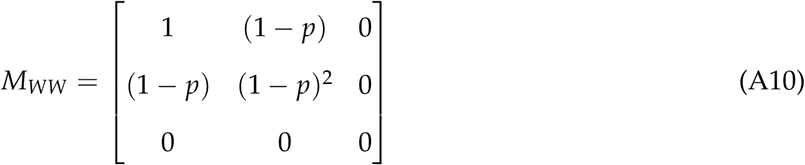

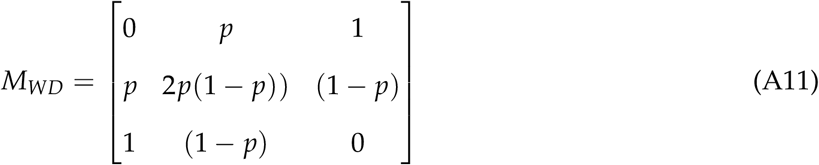

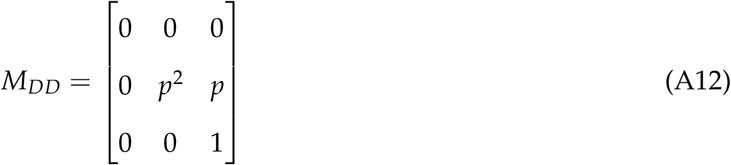

**Figure A2:**
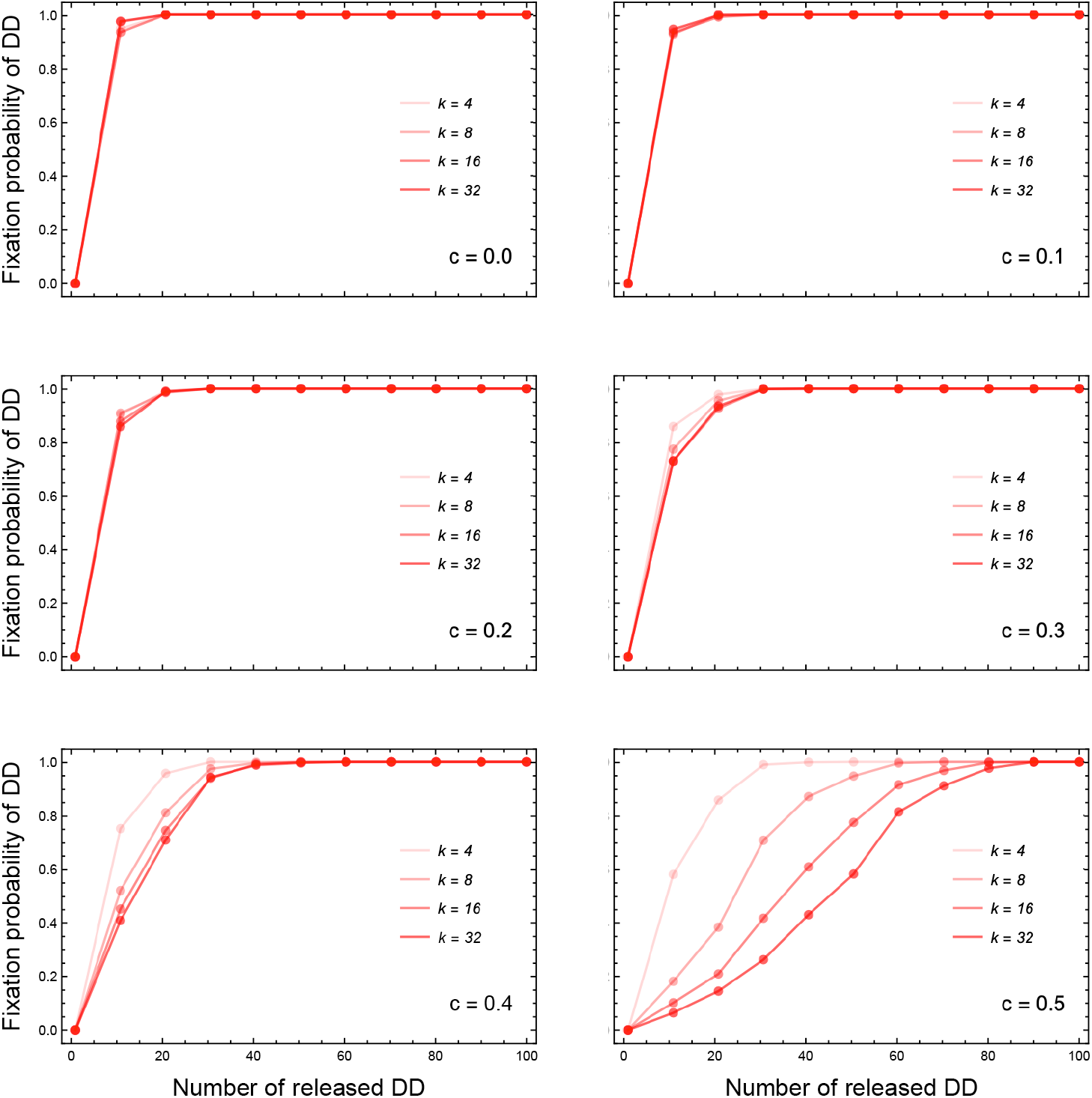
Effect of the network structure *k* on the fixation probability of *DD* for costless to costly drives (*c*). Plots show the fixation probability of drive homozygotes for networks with different average degree *k* for a variety of the number of released DD individuals. One DD individual is initially released in the population consisting only of WW. A generation consists of *n* death-birth step. We see that for a costless drive, the degree of the network does not play a major role in the fixation probability but as the cost *c* increases the more connected networks need a larger release to reach the same probability of fixation as a weakly connected network. Hence in a generation, the whole population is updated on an average. All simulations were performed for a population size of *n* = 100 and 1000 trials to estimate fixation probability. The distortion probability is set to *p* = 0.95.

## Literature Cited

Akbari, O. S., C.-H. Chen, J. M. Marshall, H. Huang, I. Antoshechkin, and B. A. Hay. 2014. Novel synthetic medea selfish genetic elements drive population replacement in drosophila; a theoretical exploration of medea-dependent population suppression. ACS synthetic biology 3:915–928.

Alphey, L. S., A. Crisanti, F. F. Randazzo, and O. S. Akbari. 2020. Opinion: Standardizing the definition of gene drive. Proceedings of the National Academy of Sciences 117:30864–30867.

Barrett, L. G., M. Legros, N. Kumaran, D. Glassop, S. Raghu, and D. M. Gardiner. 2019. Gene drives in plants: opportunities and challenges for weed control and engineered resilience. Proceedings of the Royal Society B 286:20191515.

Barton, N. H., and M. Turelli. 2011. Spatial waves of advance with bistable dynamics: cytoplasmic and genetic analogues of allee effects. The American Naturalist 178:E48–E75.

Beaghton, A., P. J. Beaghton, and A. Burt. 2017. Vector control with driving y chromosomes: modelling the evolution of resistance. Malaria journal 16:286.

Beeman, R. W., K. S. Friesen, and R. E. Denell. 1992. Maternal-effect selfish genes in flour beetles. Science 256:89–92.

Birand, A., P. Cassey, J. V. Ross, J. C. Russell, P. Thomas, and T. A. A. Prowse. 2022. Gene drives for vertebrate pest control: Realistic spatial modelling of eradication probabilities and times for island mouse populations. Molecular Ecology 31:1907–1923.

Brossard, D., P. Belluck, F. Gould, and C. D. Wirz. 2019. Promises and perils of gene drives: Navigating the communication of complex, post-normal science. Proceedings of the National Academy of Sciences of the United States of America 116:7692–7697.

Buchman, A., J. M. Marshall, D. Ostrovski, T. Yang, and O. S. Akbari. 2018. Synthetically engineered Medea gene drive system in the worldwide crop pest Drosophila suzukii. Proceedings of the National Academy of Sciences of the United States of America 115:4725–4730.

Bull, J. J. 2017. Lethal gene drive selects inbreeding. Evolution, medicine, and public health 2017:1–16.

Bull, J. J., C. H. Remien, R. Gomulkiewicz, and S. M. Krone. 2019a. Spatial structure undermines parasite suppression by gene drive cargo. PeerJ 7:e7921.

Bull, J. J., C. H. Remien, and S. M. Krone. 2019b. Gene-drive-mediated extinction is thwarted by population structure and evolution of sib mating. Evolution, medicine, and public health 2019:66–81.

Burt, A. 2003. Site-specific selfish genes as tools for the control and genetic engineering of natural populations. Proceedings of the Royal Society B: Biological Sciences 270:921–928.

Carballar-Lejarazú, R., C. Ogaugwu, T. Tushar, A. Kelsey, T. B. Pham, J. Murphy, H. Schmidt, Y. Lee, G. C. Lanzaro, and A. A. James. 2020. Next-generation gene drive for population modification of the malaria vector mosquito, anopheles gambiae. Proceedings of the National Academy of Sciences 117:22805–22814.

Champer, J., I. K. Kim, S. E. Champer, A. G. Clark, and P. W. Messer. 2021. Suppression gene drive in continuous space can result in unstable persistence of both drive and wild-type alleles. Molecular Ecology 30:1086–1101.

Champer, J., J. Liu, S. Y. Oh, R. Reeves, A. Luthra, N. Oakes, A. G. Clark, and P. W. Messer. 2018. Reducing resistance allele formation in crispr gene drive. Proceedings of the National Academy of Sciences 115:5522–5527.

Champer, J., J. Zhao, S. E. Champer, J. Liu, and P. W. Messer. 2020. Population dynamics of underdominance gene drive systems in continuous space. ACS Synthetic Biology 9:779–792.

Charlesworth, B., and D. Charlesworth. 2010. Elements of evolutionary genetics. Roberts and Company Publishers. Greenwood Village, CO.

Collins, J. P. 2018. Gene drives in our future: challenges of and opportunities for using a self-sustaining technology in pest and vector management. BMC Proceedings 12:9.

Denton, J. A., and C. S. Gokhale. 2019. Synthetic mutualism and the intervention dilemma. Life 9:15.

Deredec, A., A. Burt, and H. C. J. Godfray. 2008. The population genetics of using homing endonuclease genes in vector and pest management. Genetics 179:2013–2026.

Dhole, S., A. L. Lloyd, and F. Gould. 2020. Gene drive dynamics in natural populations: The importance of density dependence, space, and sex. Annual Review of Ecology, Evolution, and Systematics 51:505–531.

Dolezel, M. C., Lü thi, and H. Gaugitsch. 2020. Beyond limits–the pitfalls of global gene drives for environmental risk assessment in the european union. BioRisk 15:1.

Drury, D. W., A. L. Dapper, D. J. Siniard, G. E. Zentner, and M. J. Wade. 2017. Crispr/cas9 gene drives in genetically variable and nonrandomly mating wild populations. Science advances 3:e1601910.

Eckhoff, P. A., E. A. Wenger, H. C. J. Godfray, and A. Burt. 2017. Impact of mosquito gene drive on malaria elimination in a computational model with explicit spatial and temporal dynamics. Proceedings of the National Academy of Sciences 114:E255–E264.

EFSA Panel on Plant Protection Products and their Residues. 2014. Scientific opinion on good modelling practice in the context of mechanistic effect models for risk assessment of plant protection products. EFSA Journal 12:3589.

Fahse, L., P. Papastefanou, and M. Otto. 2018. Estimating acute mortality of Lepidoptera caused by the cultivation of insect-resistant Bt maize – The LepiX model. Ecological Modelling 371:50– 59.

Gantz, V. M., N. Jasinskiene, O. Tatarenkova, A. Fazekas, V. M. Macias, E. Bier, and A. A. James. 2015. Highly efficient cas9-mediated gene drive for population modification of the malaria vector mosquito anopheles stephensi. Proceedings of the National Academy of Sciences 112:E6736–E6743.

Giese, B., J. Frieß, N. Barton, P. Messer, F. Débarre, M. Schetelig, N. Windbichler, H. Meimberg, and C. Boëte. 2019. Gene drives: dynamics and regulatory matters—a report from the workshop “evaluation of spatial and temporal control of gene drives,” april 4–5, 2019, vienna. BioEssays 10:1900151.

Girardin, L., V. Calvez, and F. Débarre. 2019. Catch Me If You Can: A Spatial Model for a Brake-Driven Gene Drive Reversal. Bulletin of Mathematical Biology 81:5054–5088.

Godwin, J., M. Serr, S. K. Barnhill-Dilling, D. V. Blondel, P. R. Brown, K. Campbell, J. Delborne, A. L. Lloyd, K. P. Oh, T. A. Prowse, et al. 2019. Rodent gene drives for conservation: opportunities and data needs. Proceedings of the Royal Society B 286:20191606.

Gokhale, C. S., R. G. Reeves, and F. A. Reed. 2014. Dynamics of a combined medea-underdominant population transformation system. BMC Evolutionary Biology 14:98.

Hindersin, L., and A. Traulsen. 2015. Most undirected random graphs are amplifiers of selection for Birth-death dynamics, but suppressors of selection for death-Birth dynamics. PLoS Computational Biology 11:e1004437.

Huang, Y., A. L. Lloyd, M. Legros, and F. Gould. 2009. Gene-drive in age-structured insect populations. Evolutionary Applications 2:143–159.

Huang, Y., A. L. Lloyd, M. Legros, and F. Gould. 2011. Gene-drive into insect populations with age and spatial structure: A theoretical assessment. Evolutionary applications 4:415–428.

James, S., F. H. Collins, P. A. Welkhoff, C. Emerson, H. C. J. Godfray, M. Gottlieb, B. Greenwood, S. W. Lindsay, C. M. Mbogo, F. O. Okumu, et al. 2018. Pathway to deployment of gene drive mosquitoes as a potential biocontrol tool for elimination of malaria in sub-saharan africa: recommendations of a scientific working group. The American journal of tropical medicine and hygiene 98:1–49.

Johnson, J. A., R. Altwegg, D. M. Evans, J. G. Ewen, I. J. Gordon, N. Pettorelli, and J. K. Young. 2016. Is there a future for genome-editing technologies in conservation? Animal Conservation 19:97–101.

Karlin, S. 1978. Comparisons of positive assortative mating and sexual selection models. Theoretical population biology 14:281–312.

Kaveh, K., N. L. Komarova, and M. Kohandel. 2015. The duality of spatial death-birth and birth-death processes and limitations of the isothermal theorem. Journal of the Royal Society Open Science 2.

Leitschuh, C. M., D. Kanavy, G. A. Backus, R. X. Valdez, M. Serr, E. A. Pitts, D. Threadgill, and J. Godwin. 2018. Developing gene drive technologies to eradicate invasive rodents from islands. Journal of Responsible innovation 5:S121–S138.

Lenington, S. 1983. Social preferences for partners carrying ‘good genes’ in wild house mice. Animal Behaviour 31:325–333.

Lenington, S.. 1991. The t complex: a story of genes, behavior, and populations. Advances in the Study of Behavior 20:51–86.

Levin, S. A. 2003. Complex adaptive systems: Exploring the known, the unknown and the unknowable. Bulletin of the AMS -American Mathematical Society 40:3–19.

Lindholm, A. K., K. A. Dyer, R. C. Firman, L. Fishman, W. Forstmeier, L. Holman, H. Johannesson, U. Knief, H. Kokko, A. M. Larracuente, et al. 2016. The ecology and evolutionary dynamics of meiotic drive. Trends in ecology & evolution 31:315–326.

Lindholm, A. K., K. Musolf, A. Weidt, and B. König. 2013. Mate choice for genetic compatibility in the house mouse. Ecology and evolution 3:1231–1247.

Lindvall, M., and J. Molin. 2020. Designing for the Long Tail of Machine Learning. arXiv.

Long, K. C., L. Alphey, G. J. Annas, C. S. Bloss, K. J. Campbell, J. Champer, C.-H. Chen, A. Choudhary, G. M. Church, J. P. Collins, et al. 2020. Core commitments for field trials of gene drive organisms. Science 370:1417–1419.

Manser, A. 2015. Gene drive and sexual selection in house mice. Ph.D. thesis. University of Zurich.

Manser, A., S. J. Cornell, A. Sutter, D. V. Blondel, M. Serr, J. Godwin, and T. A. Price. 2019. Controlling invasive rodents via synthetic gene drive and the role of polyandry. Proceedings of the Royal Society B 286:20190852.

Manser, A., B. Kö nig, and A. K. Lindholm. 2020. Polyandry blocks gene drive in a wild house mouse population. Nature communications 11:1–8.

Manser, A., A. K. Lindholm, L. W. Simmons, and R. C. Firman. 2017. Sperm competition suppresses gene drive among experimentally evolving populations of house mice. Molecular ecology 26:5784–5792.

Marshall, J. M., and B. A. Hay. 2011. Inverse Medea as a Novel Gene Drive System for Local Population Replacement A Theoretical Analysis. Journal of Heredity 103:336–341.

Marshall, J. M., G. W. Pittman, A. B. Buchman, and B. A. Hay. 2011. Semele: a killer-male, rescue-female system for suppression and replacement of insect disease vector populations. Genetics 187:535–551.

Moro, D., M. Byrne, M. Kennedy, S. Campbell, and M. Tizard. 2018. Identifying knowledge gaps for gene drive research to control invasive animal species: the next crispr step. Global Ecology and Conservation 13:e00363.

Noble, C., B. Adlam, G. M. Church, K. M. Esvelt, and M. A. Nowak. 2018. Current crispr gene drive systems are likely to be highly invasive in wild populations. Elife 7:e33423.

Noble, C., J. Olejarz, K. M. Esvelt, G. M. Church, and M. A. Nowak. 2017. Evolutionary dynamics of CRISPR gene drives. Science Advances 3.

North, A., A. Burt, and H. C. J. Godfray. 2013. Modelling the spatial spread of a homing endonuclease gene in a mosquito population. Journal of Applied Ecology 50:1216–1225.

North, A. R., A. Burt, and H. C. J. Godfray. 2019. Modelling the potential of genetic control of malaria mosquitoes at national scale. BMC biology 17:1–12.

North, A. R., A. Burt, and H. C. J. Godfray. 2020. Modelling the suppression of a malaria vector using a crispr-cas9 gene drive to reduce female fertility. BMC biology 18:1–14.

O’Donald, P. 1980. Genetic models of sexual and natural selection in monogamous organisms. Heredity 44:391–415.

of Sciences Engineering, N. A., and Medicine, eds. 2016. Gene Drives on the Horizon: Advancing Science, Navigating Uncertainty, and Aligning Research with Public Values. The National Academies Press, Washington, DC.

Oye, K. A., K. Esvelt, E. Appleton, F. Catteruccia, G. Church, T. Kuiken, S. B.-Y. Lightfoot, J. McNamara, A. Smidler, and J. P. Collins. 2014. Regulating gene drives. Science 345:626–628.

Pai, A., and G. Yan. 2020. Long-term study of female multiple mating indicates direct benefits in Tribolium castaneum. Entomologia Experimentalis et Applicata 168:398–406.

Pinter-Wollmann, N., E. A. Hobson, J. E. Smith, A. J. Edelman, D. Shizuka, S. de Silva, J. S. Waters, S. D. Prager, T. Sasaki, G. Wittemyer, J. Fewell, and D. B. McDonald. 2014. The dynamics of animal social networks: analytical, conceptual, and theoretical advances. Behavioral Ecology 25:242–255.

Price, T. A., and N. Wedell. 2008. Selfish genetic elements and sexual selection: their impact on male fertility. Genetica 132:295.

Price, T. A. R., N. Windbichler, R. L. Unckless, A. Sutter, J. Runge, P. A. Ross, A. Pomiankowski, N. L. Nuckolls, C. Montchamp-Moreau, N. Mideo, O. Y. Martin, A. Manser, M. Legros, A. M. Larracuente, L. Holman, J. Godwin, N. Gemmell, C. Courret, A. Buchman, L. G. Barrett, and A. K. Lindholm. 2020. Resistance to natural and synthetic gene drive systems. Journal of Evolutionary Biology 33:1345–1360.

Prowse, T. A., F. Adikusuma, P. Cassey, P. Thomas, and J. V. Ross. 2019. A y-chromosome shredding gene drive for controlling pest vertebrate populations. Elife 8:e41873.

Prowse, T. A., P. Cassey, J. V. Ross, C. Pfitzner, T. A. Wittmann, and P. Thomas. 2017. Dodging silver bullets: good crispr gene-drive design is critical for eradicating exotic vertebrates. Proceedings of the Royal Society B: Biological Sciences 284:20170799.

Qureshi, A., A. Aldersley, B. Hollis, A. Ponlawat, and L. J. Cator. 2019. Male competition and the evolution of mating and life-history traits in experimental populations of aedes aegypti. Proceedings of the Royal Society B 286:20190591.

Rode, N. O., A. Estoup, D. Bourguet, V. Courtier-Orgogozo, and F. Débarre. 2019. Population management using gene drive: molecular design, models of spread dynamics and assessment of ecological risks. Conservation Genetics pages 1–20.

Runge, J.-N., and A. K. Lindholm. 2021. Experiments confirm a dispersive phenotype associated with a natural gene drive system. Royal Society Open Science 8:202050.

Sandler, L., and E. Novitski. 1957. Meiotic drive as an evolutionary force. The American Naturalist 91:105–110.

Simon, S., M. Otto, and M. Engelhard. 2018. Synthetic gene drive: Between continuity and novelty: Crucial differences between gene drive and genetically modified organisms require an adapted risk assessment for their use. EMBO reports 19:e45760.

Simoni, A., A. M. Hammond, A. K. Beaghton, R. Galizi, C. Taxiarchi, K. Kyrou, D. Meacci, M. Gribble, G. Morselli, A. Burt, et al. 2020. A male-biased sex-distorter gene drive for the human malaria vector anopheles gambiae. Nature biotechnology 38:1054–1060.

Singh, S. R., and B. N. Singh. 2004. Female remating in drosophila: Comparison of duration of copulation between first and second matings in six species. Current Science 86:465–470.

Sutter, A., and A. K. Lindholm. 2016. No evidence for female discrimination against male house mice carrying a selfish genetic element. Current zoology 62:675–685.

Tanaka, H., H. A. Stone, and D. R. Nelson. 2017. Spatial gene drives and pushed genetic waves. Proceedings of the National Academy of Sciences 114:8452–8457.

Unckless, R. L., A. G. Clark, and P. W. Messer. 2017. Evolution of Resistance Against CRISPR/Cas9 Gene Drive. Genetics 205:827–841.

Unckless, R. L., P. W. Messer, T. Connallon, and A. G. Clark. 2015. Modeling the manipulation of natural populations by the mutagenic chain reaction. Genetics 201:425–431.

Verma, P., R. G. Reeves, and C. S. Gokhale. 2021. A common gene drive language eases regulatory process and eco-evolutionary extensions. BMC Ecology and Evolution 21:156.

Ward, C. M., J. T. Su, Y. Huang, A. L. Lloyd, F. Gould, and B. A. Hay. 2011. Medea selfish genetic elements as tools for altering traits of wild populations: a theoretical analysis. Evolution 65:1149–1162.

Wedell, N., and T. A. R. Price. 2015. Selfish Genetic Elements and Sexual Selection, pages 165–190. Springer Netherlands.

Windbichler, N., M. Menichelli, P. A. Papathanos, S. B. Thyme, H. Li, U. Y. Ulge, B. T. Hovde, D. Baker, R. J. Monnat, A. Burt, and A. Crisanti. 2011. A synthetic homing endonuclease-based gene drive system in the human malaria mosquito. Nature 473:212–215.

Zukewich, J., V. Kurella, M. Doebeli, and C. Hauert. 2013. Consolidating birth-death and death-birth processes in structured populations. PLoS One 8:e54639.

